# Female mate choice is a reproductive isolating barrier in *Heliconius* butterflies

**DOI:** 10.1101/233270

**Authors:** Laura Southcott, Marcus R. Kronforst

## Abstract

In sexually reproducing organisms, speciation involves the evolution of reproductive isolating mechanisms that decrease gene flow. Premating reproductive isolation, often the result of mate choice, is a major obstacle to gene flow between species because it acts earlier in the life cycle than other isolating barriers. While female choice is often considered the default mode in animal species, research in the butterfly genus *Heliconius*, a frequent subject of speciation studies, has focused on male mate choice. We studied mate choice by *H. cydno* females by pairing them with either conspecific males or males of the closely related species *H. pachinus.* Significantly more intraspecific trials than interspecific trials resulted in mating. Because male courtship rates did not differ between the species when we excluded males that never courted, we attribute this difference to female choice. Females also performed more acceptance behaviours towards conspecific males. Premating isolation between these two species thus entails both male and female mate choice, and female choice may be an important factor in the origin of *Heliconius* species.

## Introduction

Speciation has produced the astounding variety of organisms that so fascinate biologists. In sexually reproducing organisms, speciation is the evolution of barriers to gene flow, creating independent lineages out of previously connected populations (Coyne and Orr 2004). Of the many barriers that can prevent interbreeding, those that occur prior to mating can exert a relatively large influence on total reproductive isolation: though hybrids may have low fertility, strong premating isolation prevents them from being formed at all (Schemske 2000; Ramsey et al. 2003). Premating barriers are especially important in cases of secondary contact or speciation with gene flow (Abbott et al. 2013).

In insects, several mechanisms can cause premating isolation: *Rhagoletis* flies mate on their host plants, so host plant preferences are a substantial barrier to hybridization (Powell et al. 2012). Damselfly species often differ in genital shape, creating a tactile cue that enables females to reject heterospecific males (McPeek et al. 2011). Songs of male *Laupala* crickets match the preferences of conspecific females (Wiley and Shaw 2010). Female, and sometimes male, preference underlies behavioural isolation between many pairs of *Drosophila* species (e.g. Coyne and Orr 1989; Noor 1995; Jennings et al. 2014), and these preferences can even be learned (Dukas and Scott 2015).

The genus *Heliconius*, containing about 45 species of Neotropical butterflies, has featured prominently in speciation research over the past three decades. They are relatively easy to rear in captivity, show geographic variation in aposematic wing colour pattern within species, and mimic both congeners and more distantly related butterfly species. *Heliconius* butterflies mate assortatively based on several cues, especially wing colour pattern (Jiggins et al. 2001; Kronforst et al. 2006b; Merrill et al. 2014). Interspecific matings produce hybrid offspring that may be sterile or more vulnerable to predators because they do not match either aposematic parental species (Naisbit et al. 2002; Merrill et al. 2012). Unlike in many taxa, male choice has been much more commonly studied than female choice in *Heliconius*, because male choice is easier to test with model females as stimuli and because male choice is the first step in mating interactions. Female choice studies have almost exclusively documented mate preference for variation (natural or experimentally-induced) in conspecific males (Finkbeiner et al. 2017; Chouteau et al. 2017; Darragh et al. 2017). However, males still regularly court heterospecific females when they have the opportunity (Merrill et al. 2011a) and researchers have generally found stronger assortative mating between species when there is the potential for female choice in the experimental design (Mérot et al. 2017). Furthermore, females are likely to bear a high cost if they hybridize, because they have low remating rates (Walters et al. 2012) and thus lower-fitness hybrids would make up most of their offspring. Therefore, female choice could facilitate speciation within the genus. The traditional focus on male mate preference in *Heliconius* research may mean we are missing a piece of the puzzle in understanding the origin and maintenance of species in this genus. Here we present an experiment to determine whether female *H. cydno* prefer males of their own species to males of the closely related *H. pachinus* (Figure 1).

**Figure 1:**
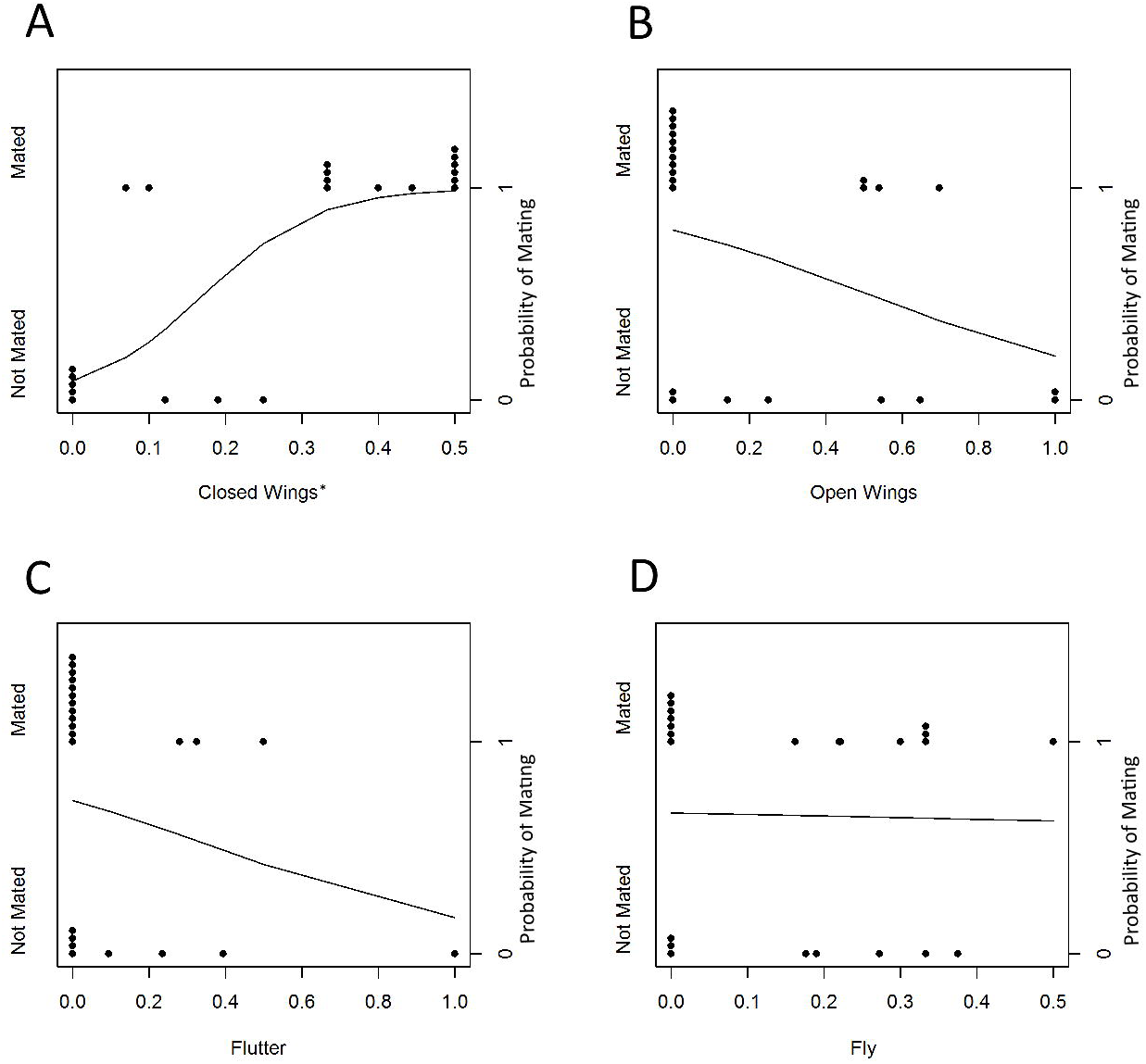
Butterfly species used in this study. A: *Heliconius cydno galanthus.* B: *Heliconius pachinus*.

## Methods

### Butterflies

*Heliconius cydno galanthus* occurs on the Caribbean coast of Central America from western Panama to southern Mexico. *Heliconius pachinus* is restricted to the Pacific coast of Costa Rica and Panama (Rosser et al. 2012). The two species diverged approximately 430,000 years ago (Kronforst et al. 2013). There is ongoing gene flow primarily from *H. pachinus* into *H. cydno* (Kronforst et al. 2013; Kronforst et al. 2006a), and hybridization is probably most prevalent around San Jose, Costa Rica, where butterflies can cross the central mountain range through a lower elevation plateau (Kronforst et al. 2007). Male mate choice contributes to reproductive isolation between the species (Kronforst et al. 2006b, Kronforst et al. 2007), but is incomplete, with 12/75 recorded matings in Kronforst et al. (2007) being interspecific.

The butterflies used in our experiments came from captive populations we established and maintained at the Smithsonian Tropical Research Institute’s insectaries in Gamboa, Panama. The captive *H. cydno* population was founded with approximately 15 wild individuals from Turrialba, Costa Rica in September 2015. The *H. pachinus* population came from eight butterflies from Reserva Forestal El Montuoso, Herrera, Panama caught in February 2016, with 15 additional wild-caught butterflies added in April 2017. Experiments took place between May 2016 and August 2017.

Adult butterflies were kept in 2.8 × 2.7 × 1.8 m cages separated by species and sex and provided with a sugar-water solution and flowers of *Lantana camara*, *Psychotria poeppigiana*, *Gurania eriantha*, *Psiguria triphylla*, and/or *Psiguria warscewiczii* daily as a pollen source. Caterpillars of both species were raised on *Passiflora platyloba* and *P. edulis* plants until pupation.

### Mate choice experiment

To test whether naive virgin female *H. cydno* prefer to mate with conspecifics, we conducted a no-choice experiment in which a female was paired with either a *H. cydno* or a *H. pachinus* male and thus given the opportunity to mate or not.

We painted females’ wings yellow to increase the probability of *H. pachinus* males approaching them. Kronforst et al. (2006b) found that *H. pachinus* males were as likely to approach wings of *H. cydno* females from a line that had a yellow forewing band introgressed from *H. melpomene* as they were to approach wings of *H. pachinus* females. We chose the simpler method of painting the forewing band to avoid the potential effects of inbreeding and *H. melpomene* genetics on female behaviour. On the day of their emergence and after their wings had fully dried, we used a Copic YG21 Anise paint pen on the dorsal surface of the forewing. This paint dries rapidly and females can fly normally within seconds of its application. Females to be paired with *H. pachinus* males had their white forewing band painted yellow, while females to be paired with *H. cydno* males had paint applied to the black part of the forewing (approximately equal area to the white band) as a control (Figure S1). Spectrophotometry indicated that painting over the black part of the wing did not substantially change its reflectance spectrum, while the yellow paint on the white band was a close approximation of the yellow pigment of *H. pachinus* and other *Heliconius* species (Figure S2). A pilot study we conducted found no difference in survival or activity levels between painted and unpainted females.

Typical courtship in adult-mating *Heliconius* begins with a male approaching a perched or flying female. The male chases a flying female, sometimes appearing to touch her, until the male breaks off pursuit or the female lands. Courtship continues with the male hovering over the perched female, possibly to waft pheromones produced by specialized androcondial scales on the wings, towards the female (Darragh et al. 2017). The male will attempt to land next to the female, facing in the same direction, and bend his abdomen towards hers to attempt to begin mating. Less commonly, males may land facing the female and touch her head with their proboscis; the function of this behaviour is unknown. Perched females may execute several behaviours during courtship. They may walk or fly away from the male; hold the wings open, preventing the male’s abdomen from reaching hers; or keep the wings closed, allowing mating to occur. She may also flutter her wings and evert her abdominal scent glands; this may be a rejection behaviour, especially in females who have previously mated. More detailed accounts of *Heliconius* courtship are given by Crane (1955, 1957), Klein and De Araújo (2010), and Jiggins (2016).

Experimental females were housed overnight in a large cage with other virgin females. Females were tested either one or two days after emergence, when they are most receptive to mating and when mating typically takes place in the wild (Jiggins 2016). A stimulus male - either *H. cydno* or *H. pachinus* at least 10 days post-emergence - was isolated in the experimental cage the day before the experiment. On the day of the experiment, the female was placed in a popup cage (30 × 30 × 30 cm) in the experimental cage for 5 minutes to acclimate. The female was then released, and both the male’s courtship attempts and the female’s responses were recorded until mating occurred or for up to 2 hours. We selected 3 male and 4 female behaviours to record based on how commonly they occur during courtship, how easily an observer can score them, and how likely they are to be correlated with the courtship’s outcome (mating or not mating). Table 1 describes the male and female behaviours recorded. Behaviours were recorded every minute, so repeated instances of the same behaviour during the same minute were not counted, but instances of two or more different behaviours during the same minute were counted. Each female and each male was used in only one experiment to ensure independence of trials.

**Table 1.** Behaviours recorded during no-choice experiment

| Behaviour | Description | Type |
| --- | --- | --- |
| **Males** |  |  |
| Chase | Male follows female closely while both are flying | Courtship |
| Hover | Male hovers over perched female | Courtship |
| Mate attempt | Male lands next to female and bends abdomen towards hers | Courtship |
| **Females** |  |  |
| Open wings | Perched female opens wings and holds them there | Rejection |
| Flutter | Perched female rapidly opens and closes wings while lifting abdomen and, usually, exposing abdominal scent glands | Rejection |
| Close wings | Perched female holds wings closed | Acceptance |
| Fly | Female flies away from male (including taking off from a perched position) | Rejection |

### Statistical analysis

We tested whether interspecific or intraspecific pairs mated more often with a chi-squared test with Yates’ continuity correction. The outcome (mating or not mating) of a no-choice trial could be attributed to male choice, female choice, or both. We tested whether males of the two species courted females equally often with aMann-Whitney U tests on the total number of courtship behaviours in a trial, the number of chases or hovers, and the number of mating attempts, excluding trials in which the male never courted. To confirm that male courtship rate did not predict the outcome of the trial, we conducted a logistic regression (GLM with a logit link function) with male species and number of courtships (the sum of all male behaviours) as independent variables and the trial outcome as the dependent variable. We examined whether females’ behaviours per male courtship predicted the outcome of the experiments using logistic regression on only the data from intraspecific trials (there were not enough interspecific matings to test whether male species interacted with these behaviours). Finally, we tested whether female behaviour rates (number of each female behaviour divided by number of male courtships) differed between inter- and intraspecific trials using Mann-Whitney U tests to determine whether females responded differently to different species of males. All analyses were performed in R (R Core Team 2013).

## Results

Intraspecific pairs mated significantly more often than interspecific pairs (Table 2; X^2^ excluding trials with no courtships: X^2^ = 9.28, df = 1, p = 0.002). *Heliconius pachinus* males were more likely to ignore the female altogether: we excluded 21 trials with *H. pachinus* males because they performed no courtship behaviours, compared to 5 such trials for *H. cydno.* We excluded trials in which the male never courted from all subsequent analyses because there is no opportunity for females to exercise choice in this context.

**Table 2.** Outcomes of no-choice experiment

|  | Mating | No mating | No courtship* |
| --- | --- | --- | --- |
| Interspecific | 4 | 21 | 21 |
| Intraspecific | 15 | 9 | 5 |

\*Trials in which males never courted the female were excluded from further analyses

The total numbers of courtship behaviours performed by male *H. cydno* and *H. pachinus* did not differ significantly (U = 338, p = 0.184; Figure S2). Numbers of hovers or chases were combined for analysis because the two behaviours were not recorded separately in some trials; the combined behaviours did not differ significantly between male species (U = 292.5, p = 0.73, Figure S2). However, *H. pachinus* males performed significantly fewer mating attempts (U = 460, p = 0.000029, Figure S2). In a logistic regression of outcome (mating or not mating) versus male species, number of courtships, and their interaction, the number of courtship attempts did not predict the outcome of the experiment (likelihood ratio tests of coefficients in a logistic regression: number of courtships p = 0.54; interaction between male species and number of courtships p = 0.57). Furthermore, comparing the full model to a reduced model with only male species as predictor, the reduced model had lower AIC (ΔAIC = 3.31) and a likelihood ratio test found that adding number of courtships did not improve the model (p = 0.71). Thus, we attribute the difference in mating rates to female preference for conspecific males rather than different intensity of male courtship once non-courting males were excluded.

In intraspecific trials, “close wings” behaviour was positively correlated with the outcome of the experiment, suggesting that wing closing indicates female acceptance of a courting male (coefficient = 14.3, SE of coefficient = 5.1, p = 0.0088). The other three behaviours were not significantly correlated with outcome, though all had negative coefficients and are considered rejection behaviours by other authors (Figure 2, Table 3, Jiggins 2016; Chouteau et al. 2017). The rates of wing opening and fluttering did not differ significantly between interspecific and intraspecific trials, although both were performed more towards *H. pachinus* males. Females closed their wings more often in intraspecific trials and flew away from the male more often in interspecific trials (Table 4, Figure 3).

**Figure 2:**
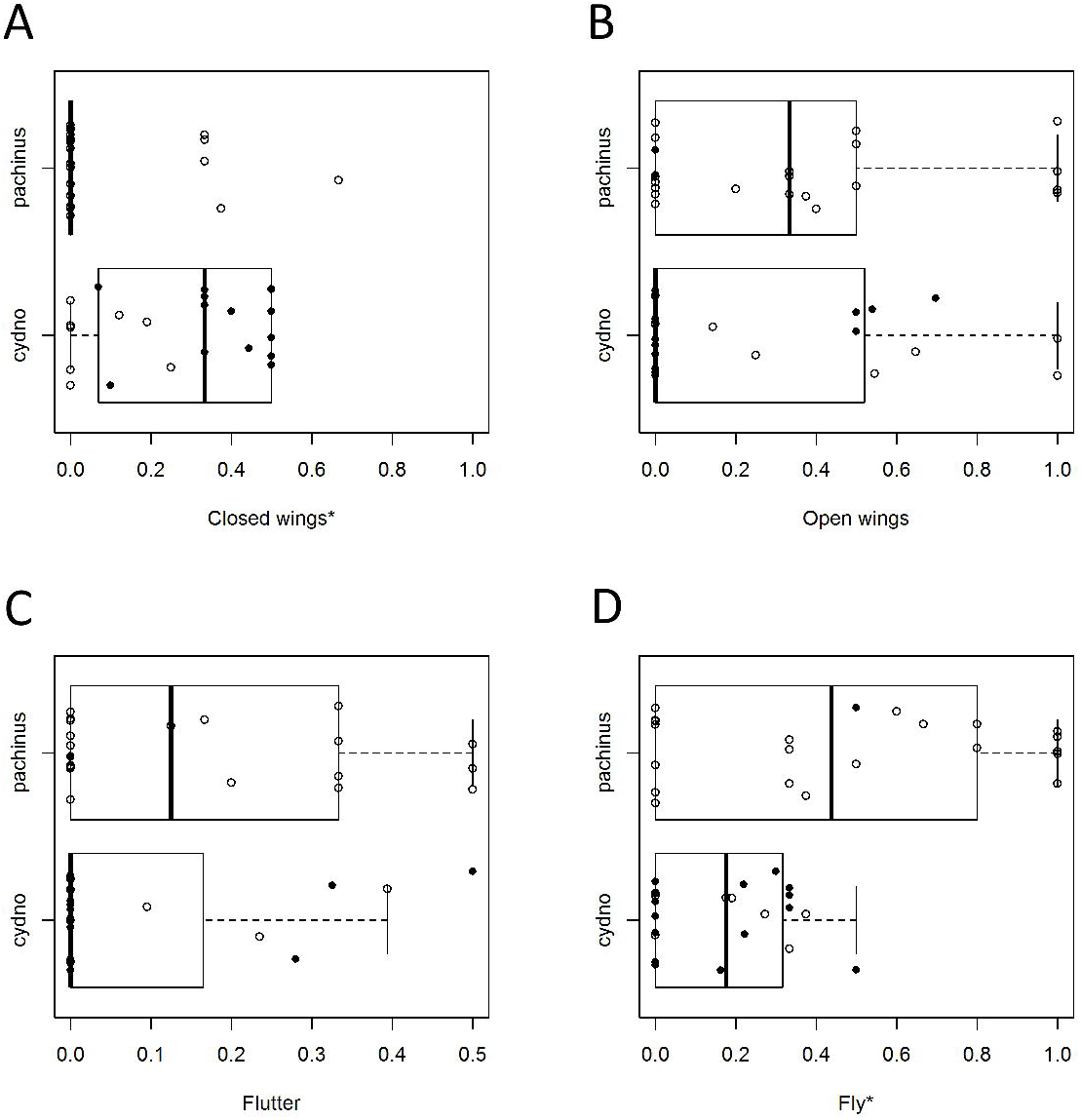
Outcome of no-choice trial and number of female behaviours per male courtship in intraspecific trials. Lines and right y-axis: Probability of mating versus female behaviour from GLMs. A: closed wings. B: open wings. C: flutter. D: fly. See Table 1 for descriptions of behaviours. Asterisks indicate behaviour that was significantly correlated with the trial’s outcome in a logistic regression.

**Table 3.** Results of logistic regressions of female behaviours per male courtship in intraspecific trials versus trial outcome (mating or no mating).

| Behaviour | Coefficient | SE | P |
| --- | --- | --- | --- |
| Closed wings | 14.3 | 5.1 | 0.0088 |
| Open wings | -2.74 | 1.5 | 0.062 |
| Flutter | -2.55 | 2.1 | 0.22 |
| Fly | -0.32 | 2.8 | 0.91 |

**Table 4.** Results of Mann-Whitney U tests comparing female behaviours per male courtship between trials with *H. cydno* (intraspecific) and *H. pachinus* (interspecific) males.

| Behaviour | U | P |
| --- | --- | --- |
| Closed wings | 369.5 | 0.001 |
| Open wings | 239 | 0.56 |
| Flutter | 200.5 | 0.12 |
| Fly | 140.5 | 0.0087 |

**Figure 3:**
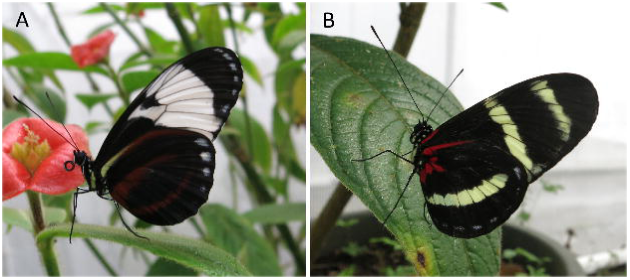
*Heliconius cydno* female behaviour rates in trials with *H. cydno* and *H. pachinus* males. A: closed wings. B: open wings. C: flutter. D: fly. Asterisks indicate behaviours whose frequency differed significantly between intraspecific and interspecific trials. Black dots: trials that ended in mating. White dots: trials that did not end in mating. Some sample sizes differ from those in Table 2 because not all behaviours were recorded in a few early trials.

## Discussion

Intraspecific no-choice trials ended in mating much more often than interspecific trials did. The lack of difference in courtship rates between species among males who courted at least once strongly suggests that female choice determined the outcome. The differences in female response behaviours (closing wings and flying away) to different species of courting males further suggests that females actively chose mates. This is the first of interspecific female choice in *Heliconius* butterflies, a model genus for speciation research with extensive evidence of male mate choice. Although male choice exists between these species (Kronforst et al. 2006b, Kronforst et al. 2007), it is weak enough that, with assistance from the manipulated female wing colour, we could observe sufficient interspecific courtships to examine females’ response to heterospecific males.

Our study adds to other attempts to document female mate preference in *Heliconius.* Recent studies have revealed intraspecific female choice in several *Heliconius* species using a variety of methods. All suggest that females exert choice during courtship based on multimodal signals, particularly vision and olfaction. In *H. erato*, females approach moving paper wings more often when they are UV reflective and have the appropriate yellow pigment (Finkbeiner et al. 2017). In the polymorphic species *H. numata*, females perform more rejection behaviours towards moving models made of dead males’ wings if those wings are of the same colour pattern morph as the female (Chouteau et al. 2017). Backcross hybrid females between *H. cydno* and *H. melpomene* are less likely to mate when they are heterozygous at the locus controlling a colour pattern element on the hindwing than when homozygous, suggesting a potential genetic component to female choice between species (Merrill et al. 2011b). In *H. timareta*, *H. erato*, and two subspecies of *H. melpomene*, females are less likely to mate with males whose pheromone-producing androconial scales have been blocked with nail varnish than with non-blocked males (Darragh et al. 2017). Similarly, perfuming conspecific males with another species’ pheromones reduced the likelihood of females mating with them (Mérot et al. 2015). Because both visual and olfactory cues differ among *Heliconius* species, these same cues could be used in interspecific female mate choice. *Heliconius cydno* and *H. pachinus*, however, have minimal differences in male pheromone composition (Schulz et al. 2007, Estrada et al. 2011), so females may be more likely to choose between these species based on visual and other non-olfactory cues.

Female choice acts within and between other butterfly species. In *Pieris occidentalis*, females prefer males of their own species over male *P. protodice*, and increasing the area of melanized spots on the forewing of *P. protodice* males increases the rate at which *P. occidentalis* mate with them (Wiernasz and Kingsolver 1992). A series of experiments revealed that *Colias philodice* females prefer conspecific males over *C. eurytheme* males, but that wing colour alone does not affect their preference (Silberglied and Taylor 1978). Females of the cryptic species *Leptidea sinapis* and *L. reali* use long courtships to distinguish between males, which court both species indiscriminately (Friberg et al. 2008). Other studies have not tested interspecific mate choice directly, but have manipulated conspecific male phenotypes. They include studies of eyespot morphology and pheromones in *Bicyclus anynana* (Robertson and Monteiro 2005; Costanzo and Monterio 2007) and of ultraviolet reflectance, iridescence, and other visual cues in *Pieris rapae* (Morehouse and Rutowski 2010), *Battus philenor* (Rutowski and Rajyaguru 2013), and *Hypolimnas bolina* (Kemp 2007), among other species.

While female choice acts in both inter- and intraspecific contexts in many butterflies, it is not always clear how much such choice contributes to total reproductive isolation. In many species, mate choice is mutual, but it is also often sequential, with males choosing whether to approach a female before the female can choose to accept or reject a male. This is certainly the case in *Heliconius*, and has long complicated efforts to detect female choice (Merrill et al. 2015). *Heliconius cydno* and *H. pachinus* are one of the younger species pairs within the genus (approximately 430 kya divergence; Kronforst et al. 2013). Compared to *H. cydno* and its next closest relative *H. melpomene*, which diverged approximately 1.4 mya (Kronforst et al. 2013), male *H. cydno* and *H. pachinus* are more likely to engage in heterospecific courtships (Merrill et al. 2011a, Mérot et al. 2017). This weaker male choice made it possible for us to induce *H. pachinus* males to court *H. cydno* females in sufficient quantities to test female choice. However, it also suggests that later in speciation between non-co-mimics female choice may decrease in relative importance among isolating barriers because male choice is strong enough that females are seldom courted by heterospecific males. Nevertheless, in young species pairs such as the one we studied, mate choice by both sexes contributes to reproductive isolation.

We have demonstrated for the first time that interspecific female mate choice is a reproductive isolating barrier between two *Heliconius* species. This finding parallels recent research showing intraspecific female choice in several *Heliconius* species and adds to the indirect case for interspecific female choice, filling a longstanding gap in the extensive literature on speciation and hybridization in this genus. Further research on the cues females use to select mates and whether they are linked to divergently selected traits is needed to understand the role of female choice in speciation.

## Acknowledgments

We thank Owen McMillan and the staff of the Smithsonian Tropical Research Institute for allowing us to use the insectary facility. We also thank Oscar Paneso, Emma Tomaszewski, Aaron Goodman, Holly Black, Amber Antonison, Lina Melo, Diego Vergara, Chantelle Doyle, and Liz Evans for capable and enthusiastic assistance with rearing butterflies, maintaining the insectaries, and performing experiments, and the rest of the STRI Gamboa community for their logistical and emotional support. We are also grateful to Trevor Price, Jill Mateo, Corrie Moreau, Claire Mérot, and two anonymous reviewers for helpful comments on this manuscript. This work was supported by the Society for the Study of Evolution (Rosemary Grant Award to LS); the University of Chicago (Center for Latin American Studies Tinker Field Research Grant and Hinds Award to LS); the Smithsonian Institution (Short-term Fellowship and CIC Graduate Fellowship to LS); National Science Foundation (grant number IOS-1452648 to MRK); and National Institutes of Health (grant number GM108626 to MRK).

