## Supplementary Materials for "Female mate choice is a reproductive isolating barrier in *Heliconius* butterflies"

### Supplemental material for: Female mate choice is a reproductive isolating barrier in *Heliconius* butterflies

September 17, 2018

#### Survival and behaviour of painted butterflies

Females were randomly assigned to two treatments: forewing bands painted as described in the main text or unpainted. We did not compare painting the forewing band vs. outside forewing band in this pilot study, as the intentation was to determine whether the handling and paint itself altered the butterflies' activity levels and survival. Both sets of females, however, had identifying marks written on their wings with an indelible marker, as had all butterflies used in the larger study.

We checked the cages where these butterflies were housed daily and collected any that had died. We recorded whether each female survived for at least one week post-treatment. Butterflies were not individually tracked for longer because the mate choice trials would only be conducted one to two days after painting. Of 14 painted butterflies, 3 died within a week of treatment; of 18 unpainted females, 1 died within a week. These proportions do not differ significantly. We note, also, that in the mate choice experiment, we did not test females that were behaving unusually (e.g. struggling to fly) on the day of the experiment (¡5 individuals).

We observed behaviour of 9 painted and 15 unpainted butterflies for 15 minutes the day after they were treated. We recorded what the female was doing at the beginning of each 30 second interval: perching, flapping wings slowly while perched, flying, walking, and feeding. Because the butterflies spent most of their time perched and the other behaviours had only a few observations per individual, we totalled all behaviours except perching and slowly flapping wings as "active" and tested whether the number of active intervals differed between treatments with a Mann-Whitney U-test. There was no significant difference in activity levels between painted and unpainted butterflies (Figure S5, $U = 48.5$, $p = 0.26$).

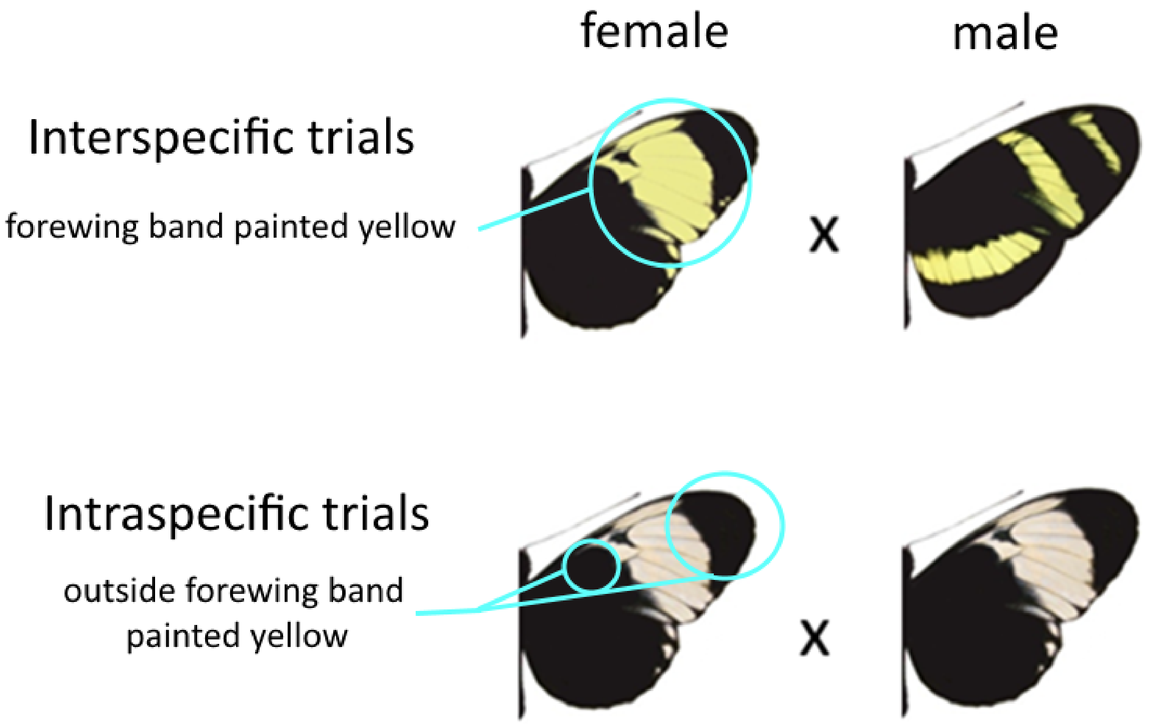


Figure S1: Experimental design: *H. cydno* females paired with *H. pachinus* males had their white forewing band painted yellow with a Copic YG21 Anise paint pen; those paired with *H. cydno* males had the black part of their forewing painted as a control.

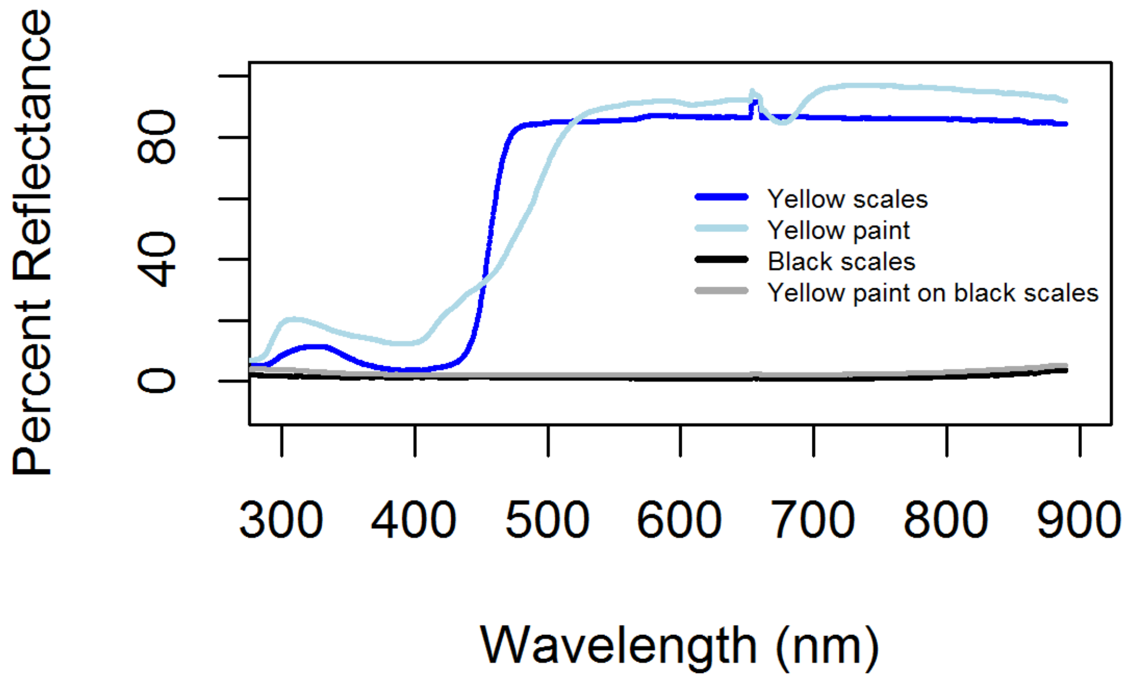


Figure S2: Reflectance spectra of painted and unpainted wing parts. A yellow morph *H. cydno alithea*, which has the same yellow pigment as *H. pachinus* (3-hydroxy-DL-kynurenine), was used as the unpainted yellow sample, and a white morph *H. c. alithea* was painted with a Copic YG21 Anise paint pen as the painted sample.

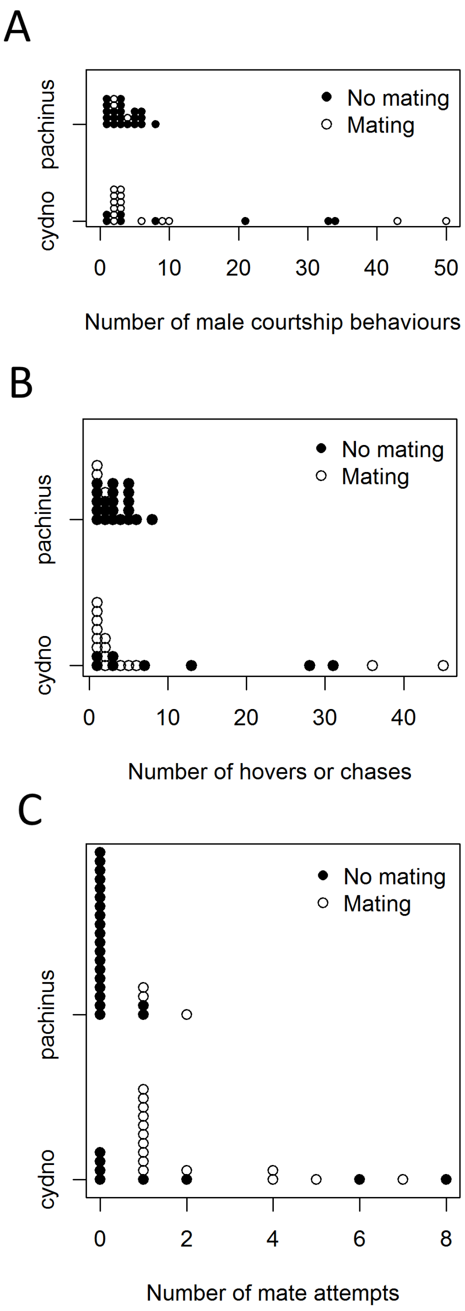


Figure S3: Courtship behaviours of *H. cydno* and *H. pachinus* males. A: Total courtship behaviours. B: Hovers or chases. C: Mate attempts. White dots: trials that ended in mating. Black dots: trials that did not end in mating.See Table 1 for descriptions of behaviours. Some sample sizes differ from those in Table 2 of the main text because not all behaviours were recorded in a few early trials.

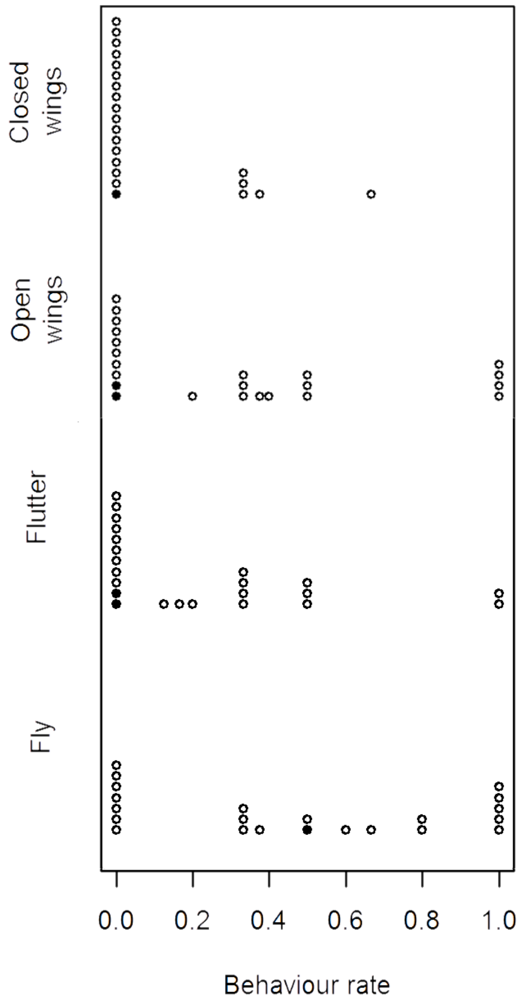


Figure S4: Rates of female behaviours (number of times female performed behaviour divided by number of male courtship behaviours) for interspecific trials (*H. cydno* females and *H. pachinus* males). Black dots: trials that ended in mating. White dots: trials that did not end in mating. Some sample sizes differ from those in Table 2 because not all behaviours were recorded in a few early trials.

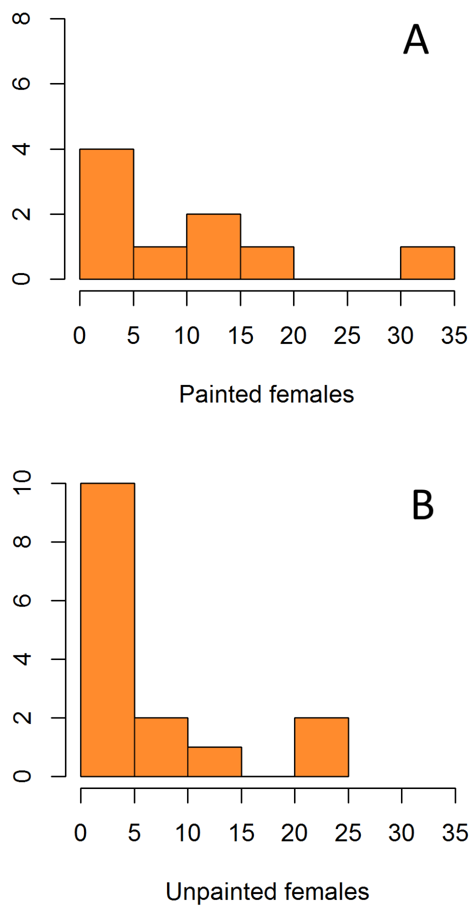


Figure S5: Number of active 30-second intervals of painted (A) and unpainted (B) butterflies.
